# Overlapping Genes and Size Constraints in Viruses - An Evolutionary Perspective

**DOI:** 10.1101/026203

**Authors:** Nadav Brandes, Michal Linial

**Affiliations:** Einstein Institute of Mathematics; Department of Biological Chemistry, Institute of Life Sciences, The Edmond J. Safra Campus, The Hebrew University of Jerusalem, Israel

**Author notes:** Corresponding author Department of Biological Chemistry, Institute of Life Sciences, The Hebrew University, The Edmond J. Safra Campus Givat Ram, Jerusalem 91904 ISRAEL; 972-54-8820035.

**Keywords:** Capsid, Icosahedral virion, ViralZone, VIPERdb, Baltimore groups

## Abstract

Viruses are the simplest replicating units, characterized by a limited number of coding genes and an exceptionally high rate of overlapping genes. We sought a unified explanation for the evolutionary constraints that govern genome sizes, gene overlapping and capsid properties. We performed an unbiased statistical analysis over the ∼100 known viral families, and came to refute widespread assumptions regarding viral evolution. We found that the volume utilization of viral capsids is often low, and greatly varies among families. Most notably, we show that the total amount of gene overlapping is tightly bounded. Although viruses expand three orders of magnitude in genome length, their absolute amount of gene overlapping almost never exceeds 1500 nucleotides, and mostly confined to <4 significant overlapping instances. Our results argue against the common theory by which gene overlapping is driven by a necessity of viruses to compress their genome. Instead, we support the notion that overlapping has a role in gene novelty and evolution exploration.

## Introduction

Viruses are the simplest biological replicating units and the most abundant ‘biological entities’ known. A great diversity is evident in their physical properties, genome size, gene contents, replication mode and infectivity. Some of the most significant properties of viruses are their small physical size and an exceptional amount of overlapping genes (OGs) relative to their genome length (Belshaw et al. 2008; Sabath et al. 2012). Most viruses have a high evolutionary rate compared to other organisms (Duffy et al. 2008; Novella et al. 2014), with that of RNA viruses 2-3 orders of magnitude higher than DNA viruses (Holland et al. 1982). The high mutation rate of RNA viruses is mostly due to the absence of a proof reading mechanism in their replicating enzymes (i.e., RNA polymerase) (Domingo 1997). It has also been shown that mutation rate is inversely correlated with genome length, not only in viruses (Duffy et al. 2008; Lynch 2010). The fast evolution of viruses is dominated by many factors, including their high mutation rate (Elena and Sanjuan 2005), large population size and fast recombination rate (Worobey and Holmes 1999). Additionally, their capacity for ‘mix and match’ during co-infection (Iyer et al. 2006; Vignuzzi et al. 2006) and for hijacking sequences from the host (Rappoport and Linial 2012) accelerate their evolutionary rate. The non-conventional evolution of many viruses leads to conflicting theories about their origin (Claverie et al. 2006; Forterre 2006; Iyer et al. 2006; Bentham et al. 2012).

Due to the inability to track the full evolutionary history of viruses, their taxonomical hierarchy is fragmented and remains debatable (Moreira and Lopez-Garcia 2009). Viruses are partitioned into 7 groups according to their genetic material and replication modes (Baltimore 1971). The two largest groups are double-stranded DNA (dsDNA) and single-stranded RNA (ssRNA+) viruses. In some families the genetic material (RNA or DNA) is segmented and composed of multiple molecules of different lengths. Different genomic segments are often packed into separate virions in the population, and a successful infection is achieved by co-infection (Moreno et al. 2014). These are collectively called segmented viruses (e.g., Brome mosaic virus, BMV) (Frank 2001).

All viruses depend heavily on their host’s translation machinery. Only a small set of proteins that fulfill the essential functions for infection are common to all viruses (Koonin 2003; Forterre 2006). These functions are restricted to: (i) recognition of the host cell, (ii) replication according to the viral group, and (iii) capsid building.

In a mature virion, the viral genome is encapsulated and protected by a capsid shell, a complex structure built of multiple (usually identical) protein subunits. The most common capsid shape is icosahedral (Zandi et al. 2006), but other structures including rod-like and irregular shapes are also known (Lidmar et al. 2003). An icosahedral capsid is composed of identical elementary protein subunits joined together in a repetitive symmetric pattern. The geometry of icosahedral solids dictates that the number of subunits can take only a fixed set of discrete values (e.g., 60, 180, etc.), determined by a property called the icosahedric triangulation (T) number (Caspar and Klug 1962). In some viruses (e.g., Simplexvirus of the family Herpesviridae), lipid layer decorated with envelope proteins surrounding the capsid shell.

A strong characteristic observed in most viruses is an abundance of overlapping open reading frames (ORFs). Many of these ORFs lack a known function (Keese and Gibbs 1992). Overlapping is a universal phenomenon, ubiquitous throughout the entire tree of life, including mammals (Veeramachaneni et al. 2004), yet only in viruses it is present in a major scale (Rogozin et al. 2002). Gene overlapping originates from various mechanisms, most notably the use of alternative start codons, ribosomal read-throughs and frame shifts (Krakauer 2000). The tendency for overlapping events is even higher in RNA viruses and in viruses with shorter genomes (Schneemann et al. 1995; Firth and Brown 2006). The codon usage characteristics was proposed to increase overlapping ORFs discovery (Pavesi et al. 2013).

Several studies have suggested various explanations for the abundance of overlapping genes (OGs) in viruses. One theory states that since viruses (especially RNA viruses) have a high mutation rate, overlapping events can increase their fitness in various ways (Krakauer 2000). For example, OGs can act as a safety mechanism by amplifying the deleterious effect of mutations occurring within them, thus quickly eliminating such mutations from the viral population (Krakauer and Plotkin 2002).

Another theory argues that overlapping has a role in gene regulation by providing a mechanism for coordinated expression. In support of this theory is the presence of OGs that are functionally related or coupled by a regulatory circuit (e.g., a feedback loop) (Krakauer 2000; Dreher and Miller 2006).

A third theory describes overlapping an effective mechanism for generating novel genes, by introducing a new reading frame on top of an existing one (Sabath et al. 2012). According to this theory, pairs of OGs are usually composed of an old well-founded gene, and a novel gene that was overprinted on top of it (Sabath et al. 2012; Pavesi et al. 2013).

The most accepted theory argues for genome compression as the driving evolutionary force (Barrell et al. 1976; Krakauer 2000; Belshaw et al. 2008; Chirico et al. 2010). Multiple arguments were raised to explain the need of viruses to have compact genomes: (i) The high mutation rate of viruses prevents them from having a long genome, as the likelihood of a deleterious mutation in each generation is length dependent (Krakauer 2000). (ii) The advantage of faster replication for infectivity. (iii) The physical size constraint imposed by the capsid’s volume (Belshaw et al. 2008). The physical size constraint is argued to be most dominant in icosahedral viruses due to the discrete nature of the T number, allowing only non-continuous changes in capsid size (Hu et al. 2008; Chirico et al. 2010). Small viruses are also argued to be subject to an even greater evolutionary pressure towards compactness, hence their high abundance of overlapping (Rancurel et al. 2009).

The motivation for this study was to systematically assess these different theories, based on an unbiased statistical analysis of the entire viral world. Currently, over 2.2 million viral proteins are archived in the UniProt public database (UniProt 2014). These proteins belong to viruses from the 7 viral groups (and additional 1% of uncharacterized proteins from metagenomic projects). We took advantage of the high-resolution structural data of some viral capsids (Shepherd et al. 2006), and the curated resource classification of viruses (Masson et al. 2013). The high quality curated database provides a non-redundant representation of reference genomes and proteomes of all known viruses.

## Results

### The landscape of overlapping genes and genome length

Although the subject of gene overlapping has already been extensively studied (e.g., (Chirico et al. 2010)), we present a revised assessment, based on the following considerations: (i) inclusion of all known viruses; (ii) unbiased sampling of the viral space based on well-curated taxa (composed of ∼400 genera within ∼100 families) as reliable representatives of the viral world; (iii) dealing only with non-trivial overlapping events (i.e., considering segments of protein-coding regions of different ORFs).

Figure 1A shows trivial and non-trivial overlapping scenarios. A trivial overlapping event is when the two genes overlap while using the same reading frame (and strand). The rest of the analysis will consider only non-trivial overlapping events (for definition, see Methods).

**Figure 1.**
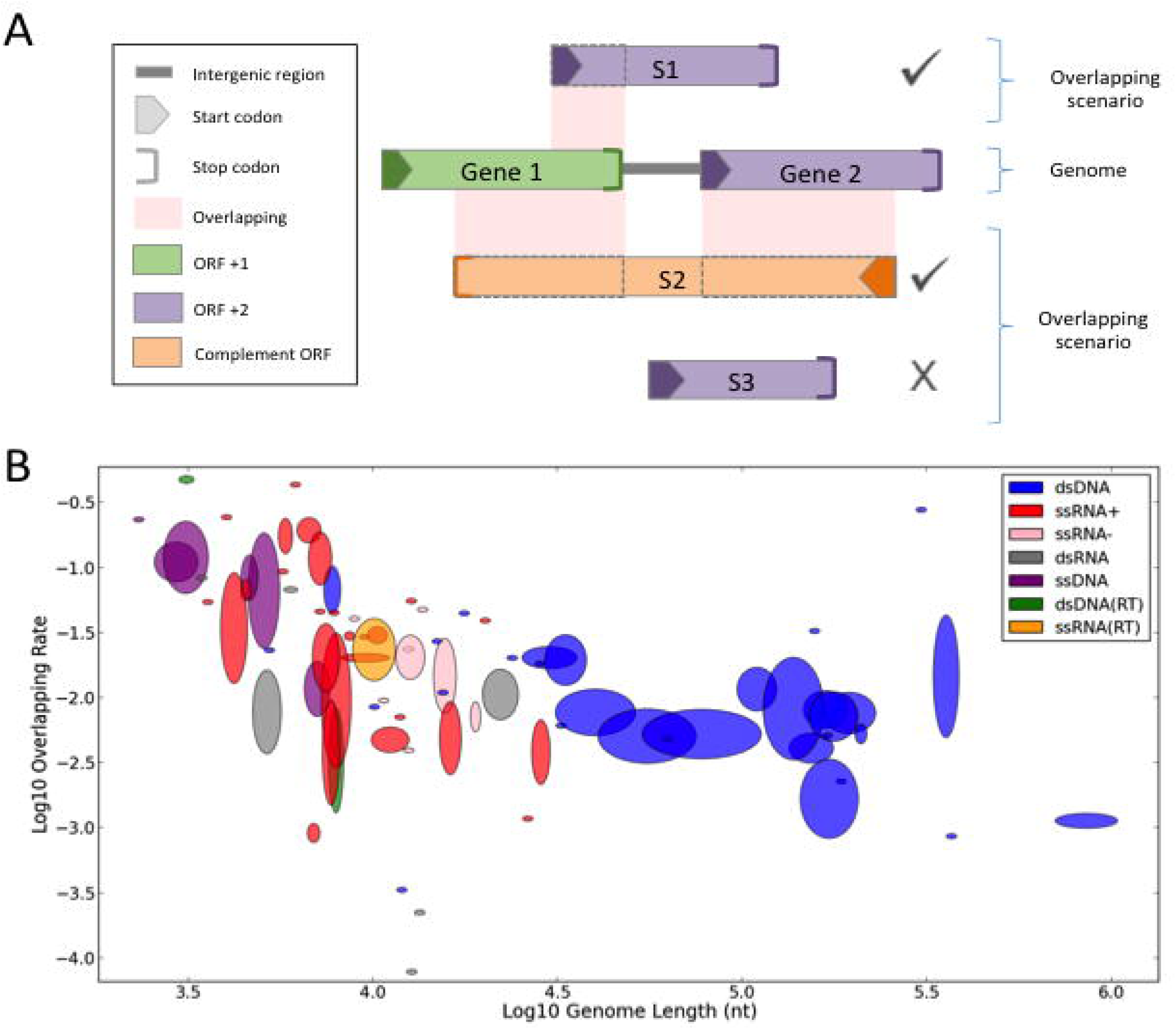
Overlapping rate is negatively correlated to genome length. **(A)** Illustration of overlapping scenarios. The definition of overlapping in this study is restricted to the presence of two genes that overlap in their coding regions while the other parts of the gene are ignored (e.g., 5’ and 3’ UTRs, intergenic). The same applies for the rare cases of genes with introns. We consider only pairs of genes that use different ORFs. It follows that the first scenario gene overlaps (marked S1) only with Gene 1, while its “overlap” with Gene 2 that share the same ORF (frame +2) is not considered (i.e., trivial overlap). The second scenario (marked S2) demonstrated that a single gene could participate in multiple overlapping events. The third scenario (marked S3) gene is not involved in any (non-trivial) overlapping event. **(B)** A scatter plot demonstrating the negative correlation between genome lengths and overlapping rate in viral families. Both axes are in log scale. 13 families without any overlapping were filtered out (to allow the use of log scale), leaving 80 families out of the complete data set of 93. The families are represented as ellipsoids, whose sizes correspond to the variance of the genera within them. The ellipsoids are colored by the partition to viral replication groups (see Methods). Spearman’s rank correlation: ρ = -0.59, p-value = 6.97 □ 10E-9.

**Error! Reference source not found.**B shows that genome length and overlapping rate (i.e., the fraction of the genome involved in overlapping; see Methods) are in a strong negative correlation, as reported before (e.g., (Belshaw et al. 2008)), meaning that smaller genomes tend have higher overlapping rates. This strong correlation (ρ = -0.59, p-value = 6.97·10E-9) remains strong when natural subsets of the viral space (e.g., single-or double-stranded viruses) are considered. In all figures, families are represented as ellipsoids, whose sizes correspond to the variance of the genera within them (see Methods).

A more direct way to measure overlapping is by absolute (rather than relative) amount. Surprisingly, the absolute amount of overlapping (measured in nucleotides, nt) remains highly bounded throughout the entire viral world (**Error! Reference source not found.**), regardless to genome length, which spans across three orders of magnitudes (∼1,500 to ∼1,000,000 nt). The absolute amount of overlapping is bounded by 1,500 nt, with only 23 of 352 genera (6.5%) and 9 of 93 families (9.7%) above this bar. When elevating the bar to 3,000 nt, only 6 of the 352 genera (1.7%) and 4 of the 93 families (4.3%) crossed it. Notably, throughout the entire spectrum of genome length, there can be found some families with a close-to-zero amount of overlapping, and other families close to the upper threshold. This is surprising, as one could have anticipated that only the viruses with high genome length will reach the upper bound.

This overlooked observation provides a stronger result than the negative correlation shown in **Error! Reference source not found.**B, which turns out to be merely a byproduct of the relative (rather than absolute) manner in which overlapping rate had been measured prior to our analysis. Specifically, when a more-or-less constant variable (absolute overlapping amount) is divided by a second variable (genome length), the division result will obviously be negatively correlated with that second variable. This is not a byproduct of using different data sets, but a direct outcome of our analysis.

**Figure 2.**
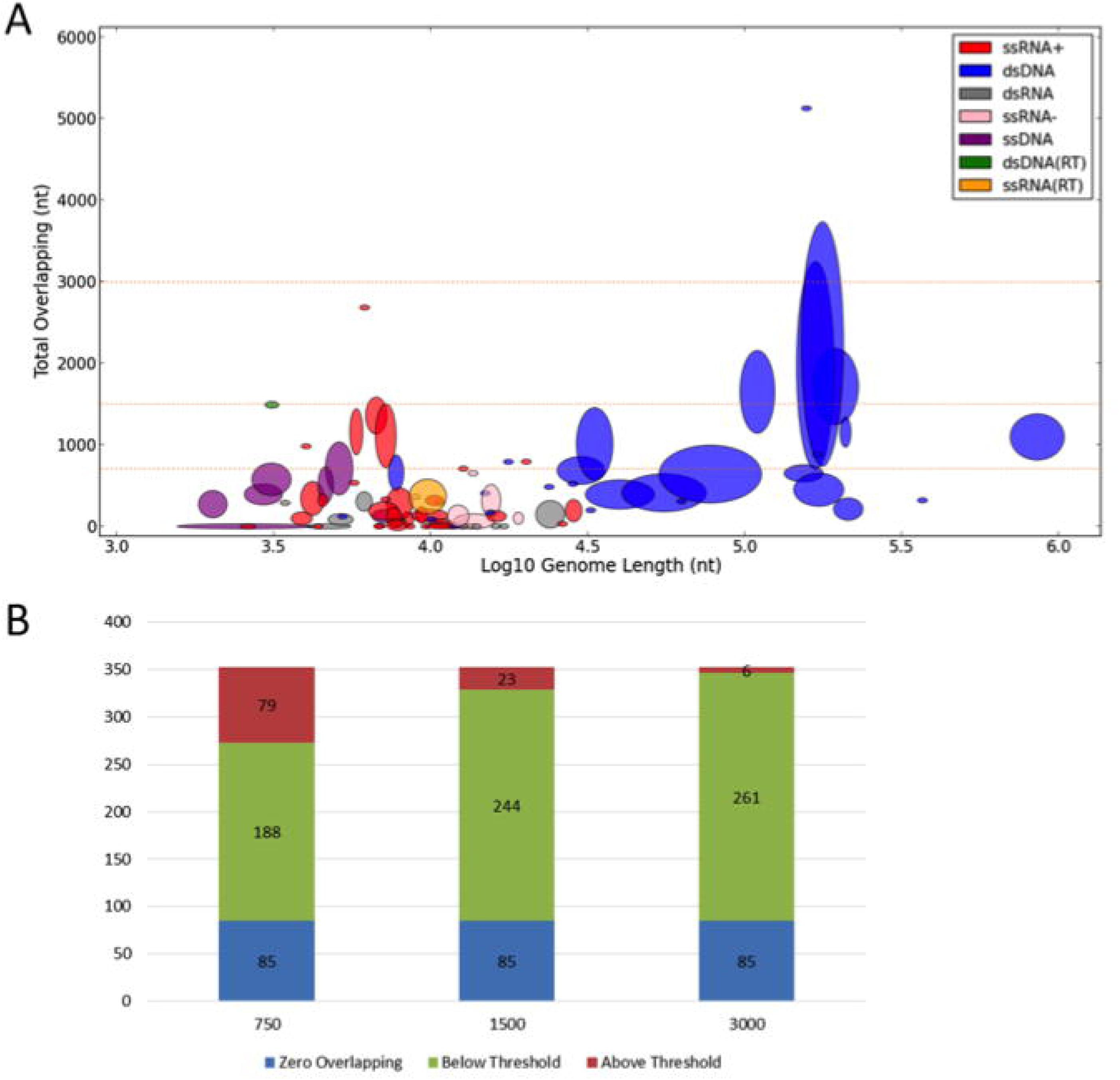
Overlapping amount is strictly bounded. **(A)** A scatter plot demonstrating the absolute number of overlapping nucleotides (nt) and genome lengths in viral families. Only the X-axis is in log scale. Throughout the entire spectrum of genome length, viral genomes have a bounded amount of nucleotides involved in overlapping. Filtered out 3 outlying families (Nimaviridae, Phycodnaviridae and Iridoviridae with 85,155, 30,798 and 7,956 overlapping nucleotides respectively), leaving 90 shown families. Spearman’s rank correlation is minimal (ρ = 0.26, p-value = 0.015). The dashed lines serve as thresholds (750, 1500 and 3000 nt) that demonstrate the bounded nature of the overlapping amount. Note that most viral families are below these bars. The different variances shown in the Y-axis when compared to Figure1B are merely a consequence of a representation of the same data (log of relative values vs. absolute values). **(B)** Of the complete data set of 352 genera, most (273, 329 and 346) have a total number of overlapping nucleotides below the chosen thresholds (750, 1500 and 3000 nt), of which 85 genera (24%) have no overlapping at all. Although the selection of thresholds is somewhat arbitrary, it can be seen that a saturation point is reached around 1500 nt.

We further tested whether our observation of a natural boundary would remain solid when counting the number of genes, rather than nucleotides, involved in overlapping. As minor overlapping events carry little constraints from evolutionary perspective (see Discussion). We considered only the subset of significantly overlapping genes (SOGs), defined by at least 300 overlapping nucleotides.

Figure 3A shows that also the number of SOGs remains highly bounded, with almost all virus families below 4 such genes, translating to less than 2 significant overlapping events. Only 3.4% of the genera and 4.3% of the families exceed this bound. Importantly, there can be found both very small and very big viruses meeting both the higher (4 genes) and lower (0 genes) bounds.

**Figure 3.**
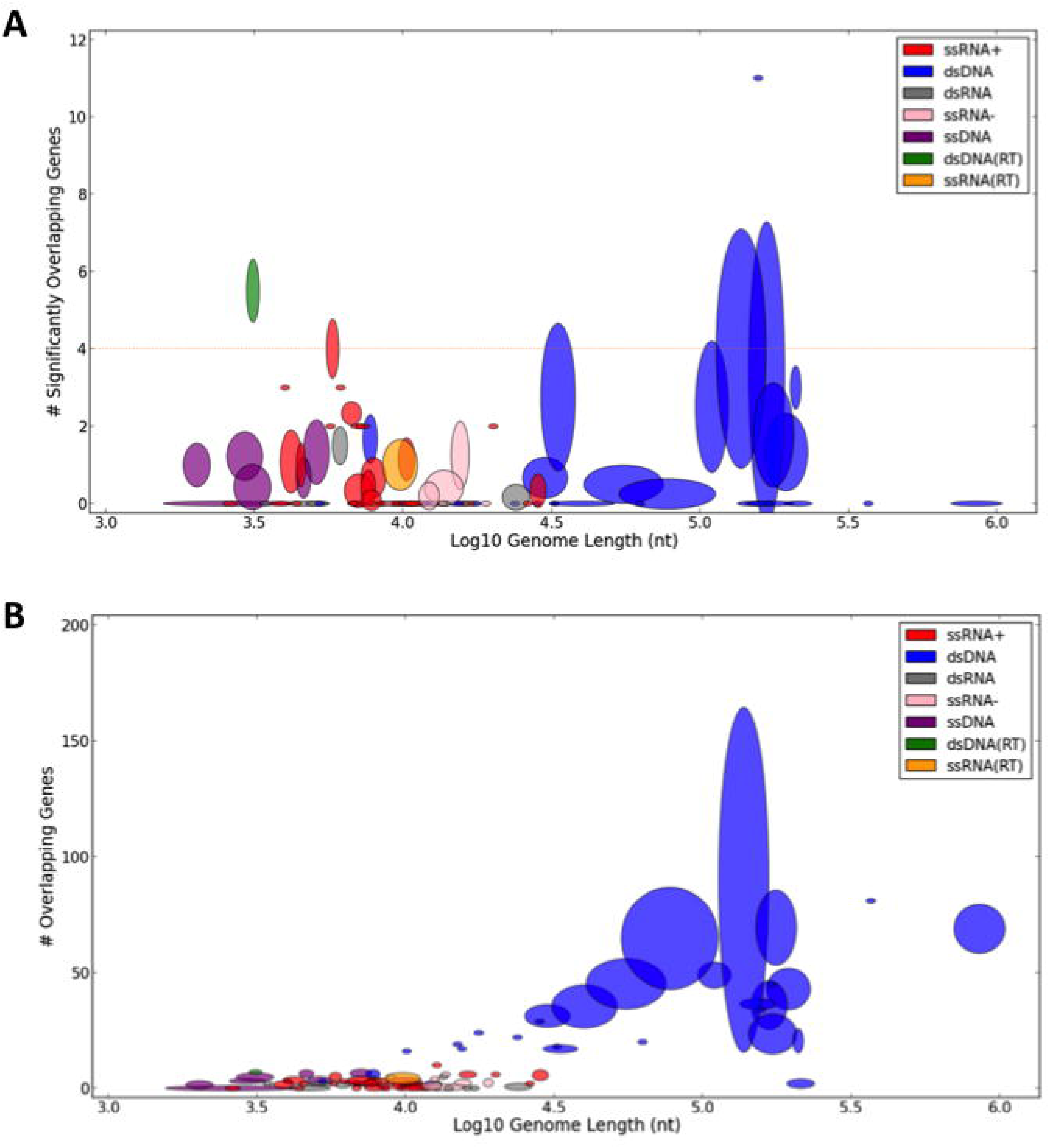
The number of significantly overlapping genes is bounded. **(A)** A scatter plot demonstrating the number of significantly overlapping genes (SOGs) with respect to genome lengths is shown for 91 of 93 viral families. Filtered out 2 outlying families (Nimaviridae and Phycodnaviridae with 141 of 532 and 50 of 505 significantly overlapping genes respectively). Only the X-axis is in log scale. Spearman’s rank correlation shows no significance (ρ = -0.08, p-value = 0.43). Most families have less than 4 significantly overlapping genes (dashed line), which account for less than 2 gene pairs. **(B)** A scatter plot demonstrating the number of all overlapping genes with no thresholds with respect to genome lengths. Only the X-axis is in log scale. Filtered out 2 outlying families (Nimaviridae and Phycodnaviridae with 489 of 532 and 283 of 505 overlapping genes respectively), leaving 91 shown families. Spearman’s rank correlation: ρ = 0.55, p-value = 1.25□10E-8.

Repeating the same analysis with varying thresholds for SOGs (50 or 100 nt, instead of 300) yields similar results (Supplemental Figure S1). However, when the threshold is eliminated altogether and all overlapping events are considered, including very minor ones (of only a few nucleotides) the total number of OGs steadily grows with genome length (Figure 3B). Since the number of SOGs remains stable, it can be deduced that only minor overlapping events that are not under evolutionary pressure become more abundant in bigger genomes (Spearman’s rank correlation: ρ = 0.55, p-value = 1.25□10E- 8).

### Overlapping is not associated with virion shape

It had been claimed that icosahedral viruses have more overlapping, as a mechanism for overcoming the unique physical constraints imposed by their capsid shape (Hu et al. 2008; Chirico et al. 2010). To test this claim, we considered the association between the physical shapes of icosahedral or non-icosahedral viruses to the phenomenon of overlapping. We revisited the viral landscape (as shown in **Error! Reference source not found.**A) and highlighted the partition between these two structural viral classes (A).

B provides a quantitative summary of these results.

It is clear that the two classes are almost indistinguishable in terms of overlapping and genome length, both showing very similar values and patterns. We conclude that, globally speaking; virion shape does not present a meaningful relation to overlapping.

**Figure 4.**
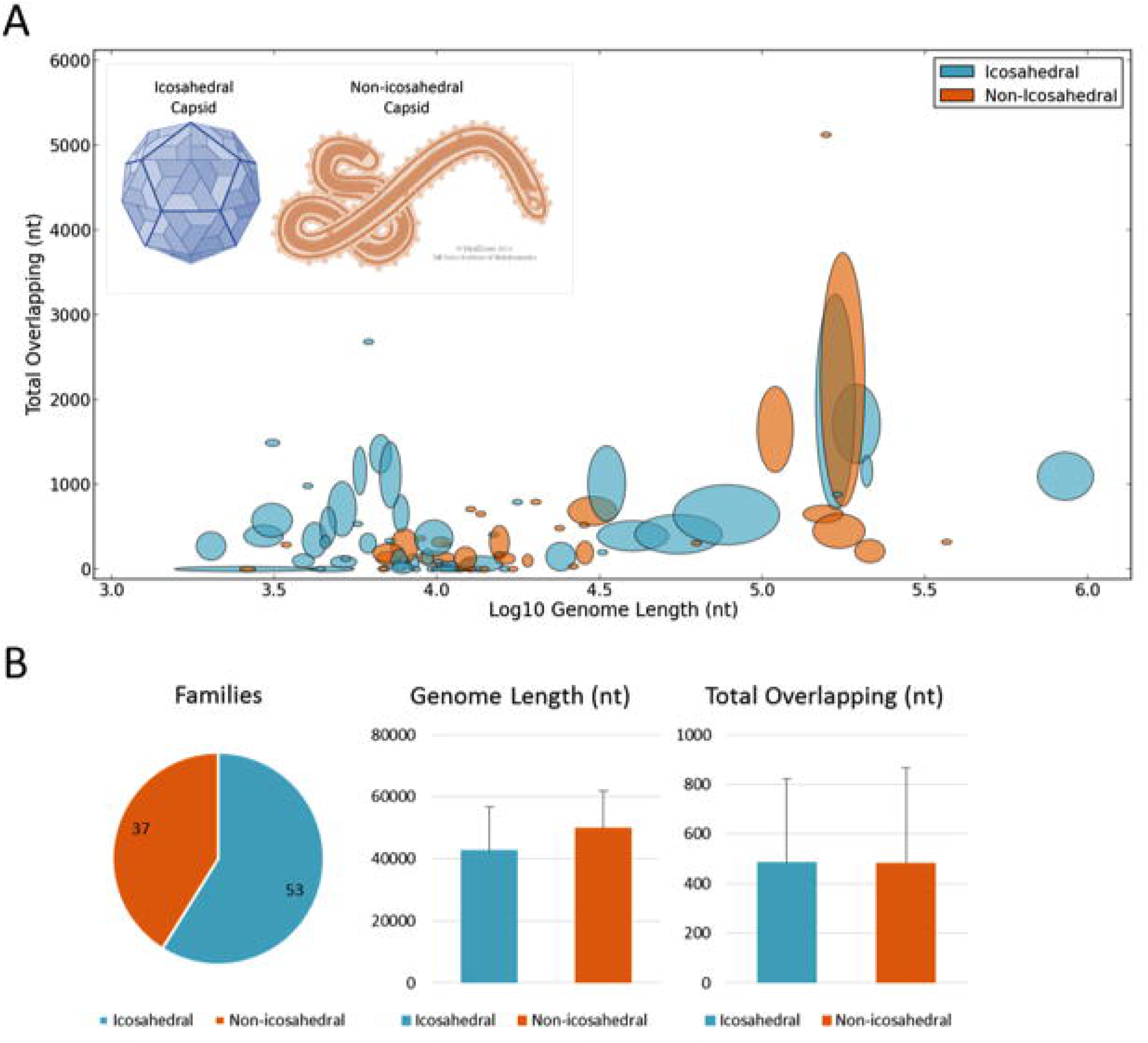
Overlapping amount and genome length are not associated with virion shape. **(A)** Showing the same analysis as in Figure 2A. The color scheme highlights the partition between icosahedral and non-icosahedral viruses. Both classes are distributed all over the space. (**B**) A quantitative summary of the 90 families in the scatter plot (37 icosahedral and 53 non-icosahedral), showing the overall statistics of the two viral classes for family resolution (see quantitative values in Supplemental Figure S3). The two classes show similar values, in terms of both average and standard deviation.

### Genome length is not constrained by capsid volume

In order to further understand whether there exist physical constraints that limit the evolution of viruses, thus driving for their exceptional rates of overlapping (**Error! Reference source not found.**B), we analyzed different aspects of capsid volumes.

We used VIPERdb (Shepherd et al. 2006), the most exhaustive resource for accurate structural data of viruses that provides detailed structural measures for icosahedral viruses. We calculated the volume usage of viruses, defined as the estimated fraction of the capsid volume occupied by the viral genome (see Methods). We found that there is no correlation (ρ = -0.17, p-value = 0.42) between the genome length and capsid volume usage among all tested icosahedral families (**Error! Reference source not found.**). The volume usage varies significantly between different viruses with no apparent pattern, and many viral families (also the very small ones) seem to be far from an optimal use of their apparent capsule space. These results remain valid also when replacing the 24 representing families with the 37 genera that compose them (Supplemental **Error! Reference source not found.**).

**Figure 5.**
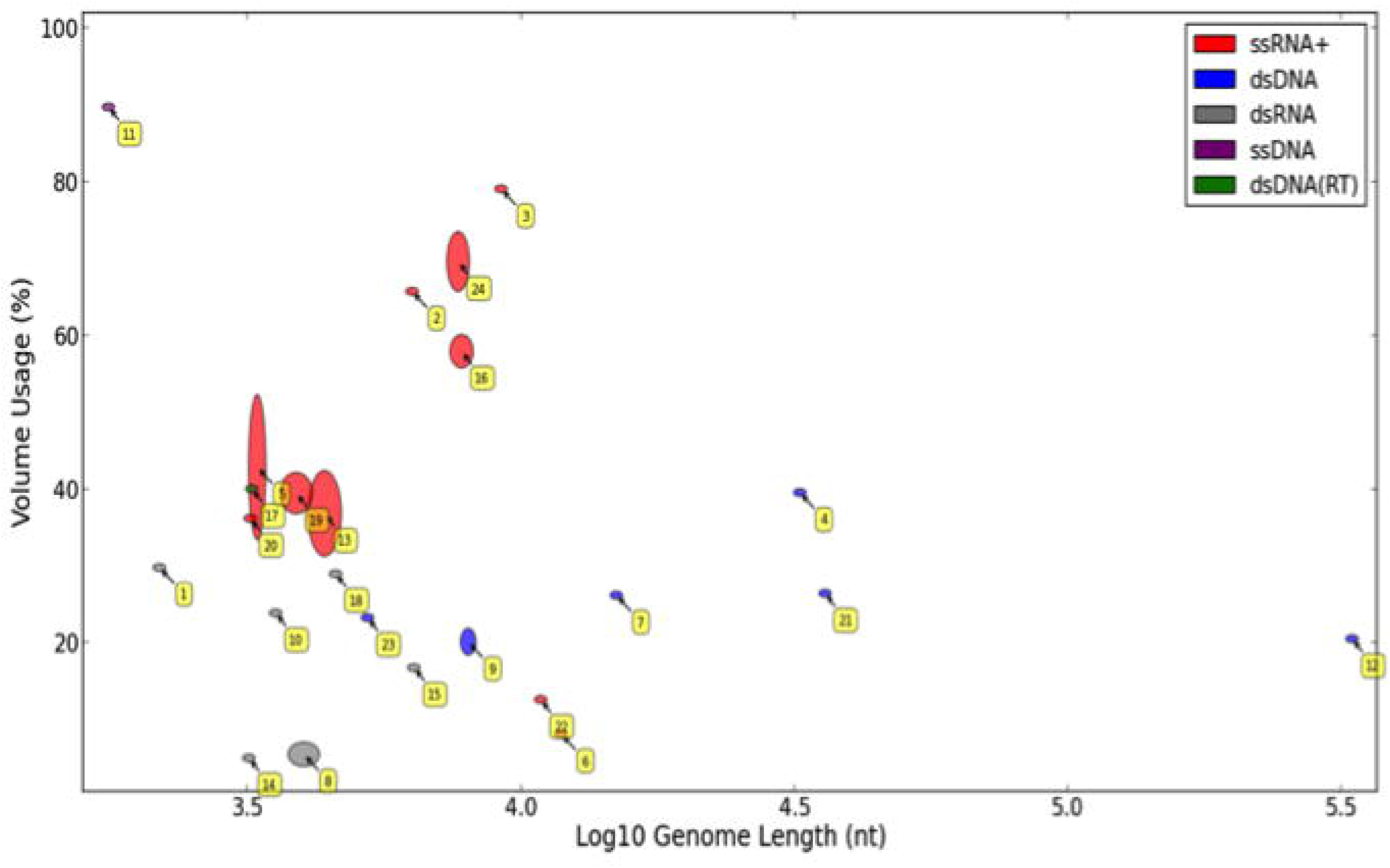
Capsid volume usage is often low and varies significantly among viral families. A scatter plot demonstrating the volume usage (in %) with respect to genome lengths is shown. Only the X-axis is in log scale. The ellipsoids are created from first calculating the volume usage percentage for each genus separately, and then drawing the families by the distributions of these values. The analysis covers all icosahedral viruses that are associated with detailed 3D information. There are 24 such icosahedral families: 1 – Partitiviridae, 2 – Tymoviridae, 3 – Dicistroviridae, 4 – Rudiviridae, 5 – Bromoviridae, 6 – Togaviridae, 7 – Tectiviridae, 8 – Reoviridae, 9 – Papillomavirida, 10 – Chrysoviridae, 11 – Circoviridae, 12 – Phycodnavirida, 13 – Tombusviridae, 14 – Birnaviridae, 15 – Cystoviridae, 16 – Caliciviridae, 17 – Hepadnaviridae, 18 – Totiviridae, 19 – Leviviridae, 20 – Nodaviridae, 21 – Adenoviridae, 22 – Flaviviridae, 23 – Polyomaviridae, 24 – Picornaviridae. Spearman’s rank correlation is not significant: ρ = -0.17, p-value = 0.42.

Table 1 provides a natural partitioning of the data presented in **Error! Reference source not found.**. Although double-stranded viruses have, in average, only half the volume usage of single-stranded viruses (24% instead of 49%), both lack a correlation between volume usage and genome length (**Error! Reference source not found.**, Table 1 and Supplemental **Error! Reference source not found.**).

**Table 1.** Volume usage in single-vs. double-stranded icosahedral families

|  | Number of Families | Average (%) | Standard deviation (%) | Spearman's rank correlation between volume usage and genome length |  |
| --- | --- | --- | --- | --- | --- |
| | | | | $\rho$ | p-value |
| Single-stranded viruses | 11 | 49 | 25 | -0.3 | 0.37 |
| Double-stranded viruses | 13 | 24 | 10 | 0 | 1 |
| All viral families | 24 | 35 | 22 | -0.17 | 0.42 |

We tested the sensitivity of the calculation towards families with segmented viruses (see Methods). We repeated the analysis by excluding all segmented viruses from (ending up with 18 families instead of 24). We observed only a minor effect of our global analysis (not shown).

Eventually, we tested the assumption that icosahedral viruses are unlikely to change their size throughout evolution (Lidmar et al. 2003; Arkhipov et al. 2006). The classification of viral genera into families, which are evolutionary related, provided us with the opportunity to measure the variation of capsid volume within families as a derivative of the extent at which viruses may adjust their capsid size with respect to their genome length.

Table 2 summarizes the variation of capsid volume inside families, with respect to capsid dimensions. It relies on atomic structural data in VIPERdb. In order to quantify variation we used the coefficient of variation (CV) statistical measure, defined as the ratio of the standard deviation σ to the mean μ, calculated individually for each family with respect to its genera. This table summarizes 40 genera in 13 families. Only families for which sufficient structural data was available are included (at least 2 genera per family). The results of this analysis demonstrate that a physical variation exists inside icosahedral families (16% and 20% on average for inner and outer volumes, respectively). Note that in many instances the differences between the inner and outer volumes are substantial. Without detailed 3D data, only the outer volume of viruses can be reliably estimated.

**Table 2.** Volume variation within icosahedral families.

| Single-stranded viruses |  |  |  |
| --- | --- | --- | --- |
| Family | Genera | Inner capsid volume CV<sup>a</sup> | Outer capsid volume CV<sup>a</sup> |
| Bromoviridae | 2 | 0.4 | 0.16 |
| Caliciviridae | 4 | 0.03 | 0.1 |
| Comoviridae | 2 | 0.11 | 0.02 |
| Leviviridae | 2 | 0.03 | 0.04 |
| Parvoviridae | 3 | 0.11 | 0.48 |
| Picornaviridae | 5 | 0.1 | 0.84 |
| Tetraviridae | 2 | 0.04 | 0.06 |
| Tombusviridae | 3 | 0.19 | 0.2 |
| **Average** | **2.88** | **0.13** | **0.24** |
| Double-stranded viruses |  |  |  |
| Papillomaviridae | 2 | 0.11 | 0.06 |
| Partitiviridae | 2 | 0.06 | 0.01 |
| Podoviridae | 4 | 0.04 | 0.03 |
| Reoviridae | 7 | 0.4 | 0.31 |
| Siphoviridae | 2 | 0.49 | 0.33 |
| **Average** | **3.4** | **0.22** | **0.15** |
| **Overall average** | **3.08** | **0.16** | **0.2** |

<sup>a</sup>CV, Coefficient of variation.

## Discussion

During our work we attempted to uncover broad unified principles that apply to most viruses. Finding global trends that apply to all viruses requires a careful unbiased approach. Obviously, our work is limited to the current coverage and classification of the viral world (Supplementary Figure S4). Due to the importance of some viruses for human health (e.g., HIV, HBV), fishery and agriculture, some viruses had been studied much more extensively than others. The outcome is an expansion in the number of reported species and genera in those well-studied families. By discussing the viruses at the family resolution, we overcome such imbalanced representation.

A major consideration in our study was to include all known viruses, using an unbiased representation. As a result, we were able to detect trends spanning three orders of magnitude in genome length, with only few outliers. Such interesting outliers (**Error! Reference source not found.**A) include the “giant viruses” Phycodnaviridae and Iridoviridae, described in the literature as very unusual in many aspects, to the extent that it was suggested to reclassify them as a new branch in the domains of life (Forterre 2010; Colson et al. 2012). Putting the 3 noticed outliers aside; the rest of the discussion will assume the validity of the trends detected across all other 90 families, discussing these results through the lens of evolution.

From evolutionary perspective, gene overlapping comes with a great price. Two functional proteins that overlap significantly (and non-trivially; **Error! Reference source not found.**A) will lead to evolutionary conflicting trends, a phenomena that was addressed as ‘constrained evolution’ (Mizokami et al. 1997). In order for a random missense mutation in an overlapping region to remain in the population, it must be beneficial for both open reading frames (or beneficial for one of them and neutral for the other). Since such an event is very unlikely, overlapping induces a great burden over the evolvement of any organism (Pavesi et al. 1997).

We confirmed the existence of a significant negative correlation between genome length and overlapping rate (ρ = -0.59, p-value = 6.97·10E-9, Figure 1B). Previous studies have interpreted this strong negative correlation as evidence in favor of the compression theory (Chirico et al. 2010) over the alternative explanations (see Introduction). However, including all families without any overlapping (13 families) the correlation become significantly weaker (ρ = -0.29, p-value = 0.0047). More critically, the observed correlation is merely a by-product of the way the overlapping is calculated (see Results). It is governed by the data representation as a *relative* value rather than by an *absolute* nucleotide counting. Instead, we found an overlooked pattern – the absolute amount of overlapping is highly bounded throughout all viruses, ranging in their length from ∼1500 to over one million nucleotides (**Error! Reference source not found.**). The compression theory does not provide an explanation to this finding. The compression theory seems especially unlikely in view of our observations in large viruses. For example, the Baculoviridae family has 4 genera, with an average of 111,260 nt genome containing 122 genes and 1,647 overlapping nucleotides. This amount of overlapping nucleotides could have theoretically been achieved by two extreme scenarios: either minor overlapping events spread over many genes, or substantial overlapping events over a small subset of genes. If compression were the main driving force for overlapping, the first strategy would be evolutionary preferred, as small overlapping events are not expected to impose significant evolutionary constraints, as opposed to substantial overlaps. We showed that minor overlaps seem not to impose an evolutionary burden as the number of OGs gradually grows with the genome length while the number of SOGs remains tightly bounded (compared Figure 3A and Figure 3B). However, it turns out that the Baculoviridae family leans more towards the second strategy. Specifically, this family has (on average) 2.5 significantly overlapping (300+ nt) genes. Moreover, the entire overlapping in this family accounts for less than 2% of its genome length, so it is unlikely that overlapping contributes significantly to compression. This argument can be generalized to most families of large viruses (**Error! Reference source not found.**A and Figure 3). Note that the total amount of overlapping does remain bounded (**Error! Reference source not found.**) by including the non-significant overlapping events, meaning that this type of overlapping events remains insignificant. Eventually, the relative perspective and the use of an inclusive definition of overlapping led to the notion that viruses have exceptional amounts of overlapping compared to other organisms (that have orders-of-magnitude larger genomes). A systematic approach had been applied to remove many of the spurious ORFs (Eberhardt et al. 2012).

Instead of the compression theory, we suggest that the observed pattern of overlapping revealed in this study is in accord with the theory of gene novelty (e.g., (Sabath et al. 2012)). According to this theory, random mutations sometimes introduce a legitimate start site on top of an existing coding gene, resulting in a new reading frame overlapping it. In fact, overlapping seems to be practically the only plausible way for viruses to increase their gene repertoire due to their compact genome organization (i.e., lack of introns and substantial intergenic regions). All other cases of gene gains must involve major genomic rearrangements or host genome contribution (e.g., gene duplication, recombination).

As the gene novelty theory predicts, it was indeed confirmed that many overlapping events involve a young (novel) gene coupled with an old well-founded partner (Sabath et al. 2012). Moreover, the signature of purifying selection was mostly found in the older of the two. For example, in the Hepatitis B virus, purifying selection was evident in only one of the paired genes (Mizokami et al. 1997). Proteins originate from OGs are characterized by being short, enriched in disordered regions, and having unusual amino acid composition (Rancurel et al. 2009). These results apply to all conformations of non-trivial overlapping. A strong argument in favor of the gene novelty theory comes from the species-specific nature of OGs (e.g., (Carter et al. 2013)). Novel OGs are generally orphans, lacking any remote homologs, unlike their older partners (Keese and Gibbs 1992).

Unlike the compression theory that could not explain the bounded amount of overlapping and other patterns observed in **Error! Reference source not found.**A and Figure 3, the theory of gene novelty provides a straightforward explanation by illustrating overlapping as a transient condition. Specifically, a significant overlapping event is not expected to last for long, due to the constant evolutionary burden imposed by it. Either one of the OGs will evolve on the expanse of the other, until it fades away, or, alternatively, they will become uncoupled (e.g., by gene duplication). Furthermore, by seeing gene novelty as the major driving force for overlapping events, it is anticipated that at any given point in time, only a small number of novel genes will be introduced to cope with the changing environment. Assuming that viruses are specified by non-redundant, indispensable gene composition, the number of gene exploration events they could tolerate simultaneously is limited. This evolutionary pressure will lead to a bounded number of OG in all viruses, and it should depend very little on their genome length, as illustrated throughout our study. Actually for large genomes of viruses (exceptions are the giant viruses), where not all the genes are indispensable we still observed a bounded number of SOGs. This observation supports the need for a limited exploration for viruses at any length, at any specific evolutionary window. The age and stability of novel ORFs is likely to be dependent on the specific viral family dynamics (e.g., (Chen and Holmes 2006).

Diverse functions provide an advantage to viruses by preoccupying the cellular systems of the host (Rappoport and Linial 2012). These functions overload the immune system (Ploegh 1998), activate ER stress (Noack et al. 2014), lead to ubiquitination and more (Hansen and Bouvier 2009; Davey et al. 2011). In other instances, the unstructured short viral peptides may cause autoimmune diseases by a molecular mimicry (Fujinami et al. 2006). An attractive rationale for the bounded number of proteins participating in OG concerns the nature of infection. When a virus enters the host cell, the cell response by activating numerous modes against the invader. Obviously, from the host cell perspective, the reaction to infection is irrespective to the size of the infected genome. According to this view, a small (bounded) number of OG suffices the condition that stimulates the host response.

A prediction of the compression theory is that viruses with smaller genomes are more densely packed (e.g., (Petrov and Harvey 2008)), presuming that meeting their maximal volume capacity prevents them from expanding their genome in the first place. Our results dispute this claim. Our reservation from this theory as the main evolutionary force does not contradict the strong tendency of viruses to be small. Viruses are indeed highly compact, in the sense of having a minimal amount of unused regulatory regions and intergenic regions (Van Etten 2003) with some exceptions (Colson et al. 2011). Furthermore, viral proteins are often shorter versions that converged toward simpler domain compositions (Rappoport and Linial 2012).

It was shown that most novel OGs are nonstructural and carry simple function (Keese and Gibbs 1992; Rancurel et al. 2009; Pavesi et al. 2013). Although degenerated, novel OGs may still be beneficial for the virus. From the perspective of information theory, overlapping does not increase the amount of information in a genome, but only partitions it among a larger set of genes, allowing more genes with less information in each. This dictates novel OGs to be poor in information, lacking complex structure and function (e.g., enzymatic), and capable of tolerating high number of mutations.

It was also claimed (Chirico et al. 2010) that icosahedral viruses have more overlapping than non-icosahedral, because the capsid size of the former is less flexible and unable to grow continuously, consequently these viruses are not capable of increasing their genome length. Our results dispute these claims. First, the pattern of overlapping and genome length is similar in both icosahedral and non-icosahedral viruses (, Supplemental Table S4). Moreover, if there is any difference in the variance of genome length inside families between these two classes, icosahedral viruses are in fact the ones with a slightly higher variance, suggesting that they are indeed capable of increasing or decreasing their genome length. It may still be that the higher variance observed in icosahedral families is merely a bias caused by the fact that an icosahedral family has more recorded genera on average (4.6 instead of 2.7).

Are icosahedral capsids unable to continuously change along evolution? Although changing the T number would result a major change in the capsid size, it might indeed be possible to slightly change the size of each subunit composing the capsid. Indeed, a variance in both the inner and outer capsid volumes exists among the genera of icosahedral families (Table S2). Our structural results undermine the common claim that the alleged compression requirement of viruses is driven by physical size constraints imposed by a limited space in their capsid. **Error! Reference source not found.** shows a great variance in volume usage among families (distributed with no apparent pattern), suggesting that physical space is probably not a significant constraint for viral evolution, as many viruses, even small ones, use only a small fraction of the volume available for them. The observation that the volume usage of single-stranded viruses is significantly higher than that of double-stranded (49% vs. 24% on average, Table 1) remains unexplained. However, in some families the viruses are packed with additional proteins that are essential for the infectivity (e.g., Vif protein in HIV (Liu et al. 1995)). Others carry replication (polymerase) or packing proteins. The volume usage estimation ignore the contribution of host derive particle-associated proteins (Belshaw et al. 2008). In most instances these proteins occupies a minor fraction of the inner volume. Eventually, there are different mechanisms for packing viral genomes inside a capsid (Snijder et al. 2013). In bacteriophages, the packing of the dsDNA is essential for a successful ejection during infection. On the other hand, effectible packing and compressing single-stranded genomes (RNA and DNA) is based on electrostatic interaction of the capsid proteins with the nucleic acids negative charges (Belyi and Muthukumar 2006).

One would quickly find out that it is a lot easier to make hypotheses about the entire viral world rather than proving them. This complex behavior of volume usage raises concerns about the interpretation of a recently reported study showing a strong linear correlation between the logarithm of viral genome lengths to the logarithm of their capsid volumes (Cui et al. 2014). It was originally interpreted that a strong polynomial relationship exists between these two variables (since log *y* ≈ as *A* log *x* + *B* suggests *y* ≈ *e^B^x^A^*) and that “virion sizes in nature can be broadly predicted from genome sequence data alone”. Although we obtain a similar linear correlation with Spearman’s rank correlation: ρ = 0.83, p-value = 4.15□10E-7 (Supplemental Figure S3). Our analysis does not support a polynomial model. We have demonstrated a great variation in volume usage, with most viruses in the range of 20-80% (**Error! Reference source not found.**), meaning that predicting the capsid size from genome length cannot be accurate. Indeed, the suggested polynomial model contains errors of up to an order of magnitude (Cui et al. 2014). Furthermore, this polynomial model is not robust to natural partitioning of the data. For example, the results of the linear regression change dramatically from a coefficient of 0.9 in double-stranded to 1.58 in single-stranded viruses (where the coefficient for both is 1.13). Obviously, these give very different polynomial models, y= *C*_1_ *x*^O.9^ vs. y= *C*_2_ *x*^l.58^, suggesting that over-fitting is involved.

A general conclusion from our study is that viruses are much more flexible than often claimed, as witnessed in their size patterns. Drawing a general model and rigid constraints for the sizes of all viruses seems infeasible when dealing with such a diverse world. Nevertheless, we were able to show that a universal boundary on overlapping exists which applies to viruses from all replication groups, viruses having 1000 folds difference in their genome length, and viruses that are either icosahedral or having any other shape. We conclude that this overlooked trend is best explained through the lens of dynamic changes with respect to changing environments (e.g., their host), and the concepts of infection that is shared by all viruses, irrespectively to their size.

As our results are relying on a statistical analysis, we do not expect them to apply to every single family, nor to all possible subsets of the data. It is likely that special viral taxa do not follow some of the general trends we have found. We share the data and the computational code to assist researchers to further use this data (including many attributes). Understanding the driving forces and constraints that govern viral evolution becomes highly relevant in view of epidemic episodes and outbreaks in recent years (e.g., (Smith et al. 2009)). The task of developing sustainable antiviral treatment strategies and sophisticated viral-based delivery systems heavily depends on it (Kay et al. 2001; Ghedin et al. 2005).

## Methods

### Data and Resources

We used two main data sources: ViralZone ((Masson et al. 2013); http://viralzone.expasy.org/) and VIPERdb ((Shepherd et al. 2006); http://viperdb.scripps.edu/). ViralZone has been used for a taxonomical categorization of the viral world as presented by the International Committee on Taxonomy of Viruses (ICTV). All species viruses are classifies to replication groups, families, genera and species (see Supplementary Figure S4). It is linked to genomic data, through reference genomes from NCBI (Pruitt et al. 2012). In addition, when structural data could not be found at VIPERdb for certain viral families, we also searched inside ViralZone pages for information about their icosahedral T numbers. Specifically, the T number information has been used to distinguish between icosahedral and non-icosahedral families. We assumed that a family is icosahedral if and only if it appears in VIPERdb or has a T number in ViralZone.

From VIPERdb we extracted capsid structural data, specifically the radiuses used for all the volume analyses. VIPERdb also classify the records by families and genera. We used this classification in order to match between ViralZone and VIPERdb records, providing us with both genomic and structural data for the common genera that appear in both resources.

VIPERdb document all the solved structures of icosahedral capsids (linked to its PDB record (Holmes 2011)). Each genus might have dozens of different records, several of the records represent mutated version of capsid proteins. In this study we were obviously interested in naturally occurring infective viruses. We thus combined all VIPERdb records sharing the same genus, ending up with a single record for each genus. When multiple values were available for a certain genus, we took the maximal value to represent the genus. Using this protocol we overcome the cases in which the capsid subunits (often a mutant protein) collapse inwards to form a shape that is incompatible with a capsid that is capable of containing its genome (Wright 2014). For example, the Mastadenovirus genus (of the family Adenoviridae in the dsDNA group) has 10 records in VIPERdb, with inner radius values ranging from 41Å to 326Å. These radii represent an artifact of an empty capsid and a natural infective virion, respectively. The inner radius of the infective virion ranges from 311-326Å. We merged these 10 records into a single record representing the Mastadenovirus genus, whose inner radius was set to be 326Å. A similar pattern occurs in most genera.

In order to retain maximal objectivity, it was crucial that the entire process of data extraction would be automatic, without any local decisions being made for specific records. For this reason all the data was extracted from the different databases (ViralZone, VIPERdb and NCBI) using unbiased downloading protocols and the analysis was performed on the entire set of records. Note that when we removed outliers it was merely for the sake of figure visibility. We clearly indicate the removed outliers in the figures’ caption and in the statistical analysis.

After extracting the data, we ended up with full taxonomic and genomic information of 352 genera in 93 families taken from ViralZone and its NCBI links. This number is slightly lower than the 420 genera reported in ViralZone (April 2015). The missing genera did not have a complete reference species. We processed 419 VIPERdb records, which we grouped into 68 genera in 37 families. For 43 genera in 28 families we had records from both ViralZone and VIPERdb. The set of 28 families were applied for volume analyses (see “volume calculations” section).

All the data we extracted are available in the Supplemental Tables S1-S3. These files contain additional fields and properties that can be useful for a follow up research. We share our Python code, which contains a handy framework for analyzing this data (Supplemental file S1).

### Taxonomy and representative selection

Different families might dramatically vary in the number of recognized genera they cluster together (e.g., the Picornaviridae family has 23 reference genera in ViralZone, while many other families have only 1). In this study we sought to conduct an unbiased statistical analysis. Thus, we conduct this study mostly at the family resolution, giving an equal weight to all viral families, regardless of the number of genera and species they might have.

For each variable involved in the analysis, we took the family’s value to be the average among all of its genera. When calculating Spearman’s rank correlation, for example, the samples used for the statistical test were actually the average values of each family. Yet, in order to also show the variety that might exist within families, we drew each family as an ellipse. The ellipse’s center corresponds to the average value of the family’s genera, and its width and height correspond to the standard deviations of its genera with respect to each of the two studied variables.

Throughout this study we ignored the variation within genera, taking the value of each genus to be the maximum among its species, doing so for each property separately. For example, the genome length of a genus was determined by its species with the maximal genome length.

### Overlapping measurements

We defined the amount of overlapping in a genome to be the number of nucleotides (nt) involved in a non-trivial gene overlapping events. A trivial overlapping event is when the two genes overlap but the same reading frame (and strand) is used. The majority of overlapping instances in viruses are trivial, where the end product is an extended version of the same protein with alternative start or stop sites (obviously, this leads to more than one protein with the same amino-acid sequence coded by the overlap region). Trivial overlapping lacks all the interesting evolutionary implications, hence we removed it of the analysis. Also, whenever referring to genes, only protein-coding regions are considered. It follows that overlapping amount, which is given in nucleotides, can immediately be translated to amino acids (i.e., 3nt to 1aa).

Overlapping rate is defined as the relative part of the genome involved in overlapping (i.e., the amount of overlapping divided by the genome length). We defined an overlapping event to be significant (coined SOG), if it involves at least 300 nt from both OGs. As we have demonstrated, our results are not sensitive to this exact threshold, but having a threshold is crucial (see Discussion). Recall that every overlapping event involves at least two genes, so when talking about the number of genes in a genome involved in overlapping, the number of overlapping events is usually only half that number.

### Volume calculations

Most volume analyses were limited to the 28 viral families for which we had both high quality genomic data from NCBI (linked from ViralZone) and capsid structural data (from VIPERdb). We defined the volume usage of a virus to be the ratio of its genome volume to the volume of its capsid. Some genera resulted an apparent volume usage that exceeds 100%. These are capsid shapes that resulted from artificial capsid shape of mutated proteins. In vitro assemble of capsid without its genome is often resulted in a collapsed shape. After applying this filter, we ended up with 24 families that had at least one proper genus.

Icosahedral solids are roughly spherical, so we calculated their volumes by the formula of a ball’s volume: 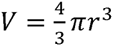, where r is the capsid’s inner radius, as provided by VIPERdb. Genomic volumes were calculated assuming that double-stranded DNA (or RNA) molecules are roughly cylindrical with a ∼20Å diameter and a distance of ∼3.4Å between adjacent nucleotides in the backbone (Arsuaga et al. 2002), yielding 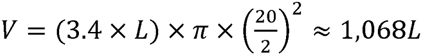, where L is the genome length (in nt). For single-stranded genomes we took half that volume (i.e., v ≈ 534L). This calculation ignores higher-order conformations of the genomic material, making it only a lower bound to the true genomic volume. Despite the limitation in calculating the usage of the capsid volume, we would still expect to see a uniform volume usage for the different families if the idea that viruses utilize their available space was correct. Our results would still suggest that many viruses do not fully utilize their available capsid volume.

Another complication in calculating genome volume arises from segmented viruses (see background). It was shown that different particles most likely have only a subset of the segments (Frank 2001). We calculated genome volume based on the length of the longest segment.

## Acknowledgements

We thank Nati Linial for useful discussion and critical comments. We thank Liran Carmel and Kerem Wainer for critical reading of the manuscript. The research was supported in part by EU FRVII Excelerate grant of Elixir.

## References

Arkhipov A, Freddolino PL, Schulten K. 2006. Stability and dynamics of virus capsids described by coarse-grained modeling. Structure 14(12): 1767–1777.

Arsuaga J, Tan RK, Vazquez M, Sumners DW, Harvey SC. 2002. Investigation of viral DNA packaging using molecular mechanics models. Biophysical chemistry101-102: 475–484.

Baltimore D. 1971. Expression of animal virus genomes. Bacteriological reviews 35(3): 235–241.

Barrell BG, Air GM, Hutchison CA, 3rd. 1976. Overlapping genes in bacteriophage phiX174. Nature 264(5581): 34–41.

Belshaw R, Gardner A, Rambaut A, Pybus OG. 2008. Pacing a small cage: mutation and RNA viruses. Trends in ecology & evolution 23(4): 188–193.

Belyi VA, Muthukumar M. 2006. Electrostatic origin of the genome packing in viruses. Proceedings of the National Academy of Sciences of the United States of America 103(46): 17174–17178.

Bentham M, Holmes K, Forrest S, Rowlands DJ, Stonehouse NJ. 2012. Formation of higher-order foot-and-mouth disease virus 3D(pol) complexes is dependent on elongation activity. Journal of virology 86(4): 2371–2374.

Carter JJ, Daugherty MD, Qi X, Bheda-Malge A, Wipf GC, Robinson K, Roman A, Malik HS, Galloway DA. 2013. Identification of an overprinting gene in Merkel cell polyomavirus provides evolutionary insight into the birth of viral genes. Proceedings of the National Academy of Sciences of the United States of America 110(31): 12744–12749.

Caspar DL, Klug A. 1962. Physical principles in the construction of regular viruses. Cold Spring Harbor symposia on quantitative biology 27: 1–24.

Chen R, Holmes EC. 2006. Avian influenza virus exhibits rapid evolutionary dynamics. Molecular biology and evolution 23(12): 2336–2341.

Chirico N, Vianelli A, Belshaw R. 2010. Why genes overlap in viruses. Proceedings Biological sciences / The Royal Society 277(1701): 3809–3817.

Claverie JM, Ogata H, Audic S, Abergel C, Suhre K, Fournier PE. 2006. Mimivirus and the emerging concept of “giant”. tvirus. Virus research 117(1): 133–144.

Colson P, de Lamballerie X, Fournous G, Raoult D. 2012. Reclassification of giant viruses composing a fourth domain of life in the new order Megavirales. Intervirology 55(5): 321–332.

Colson P, Yutin N, Shabalina SA, Robert C, Fournous G, La Scola B, Raoult D, Koonin EV. 2011. Viruses with more than 1,000 genes: Mamavirus, a new Acanthamoeba polyphaga mimivirus strain, and reannotation of Mimivirus genes. Genome biology and evolution 3: 737–742.

Cui J, Schlub TE, Holmes EC. 2014. An allometric relationship between the genome length and virion volume of viruses. Journal of virology 88(11): 6403–6410.

Davey NE, Trave G, Gibson TJ. 2011. How viruses hijack cell regulation. Trends in biochemical sciences 36(3): 159–169.

Domingo E. 1997. Rapid evolution of viral RNA genomes. The Journal of nutrition 127(5 Suppl): 958S– 961S.

Dreher TW, Miller WA. 2006. Translational control in positive strand RNA plant viruses. Virology 344(1): 185–197.

Duffy S, Shackelton LA, Holmes EC. 2008. Rates of evolutionary change in viruses: patterns and determinants. Nature reviews Genetics 9(4): 267–276.

Eberhardt RY, Haft DH, Punta M, Martin M, O’Donovan C, Bateman A. 2012. AntiFam: a tool to help identify spurious ORFs in protein annotation. Database : the journal of biological databases and curation 2012. bas003.

Elena SF, Sanjuan R. 2005. Adaptive value of high mutation rates of RNA viruses: separating causes from consequences. Journal of virology 79(18): 11555–11558.

Firth AE, Brown CM. 2006. Detecting overlapping coding sequences in virus genomes. BMC bioinformatics 7: 75.

Forterre P. 2006. The origin of viruses and their possible roles in major evolutionary transitions. Virus research 117(1): 5–16.

Forterre P.. 2010. Giant viruses: conflicts in revisiting the virus concept. Intervirology 53(5): 362–378.

Frank SA. 2001. Multiplicity of infection and the evolution of hybrid incompatibility in segmented viruses. Heredity 87(Pt 5): 522–529.

Fujinami RS, von Herrath MG, Christen U, Whitton JL. 2006. Molecular mimicry, bystander activation, or viral persistence: infections and autoimmune disease. Clinical microbiology reviews 19(1): 80–94.

Ghedin E, Sengamalay NA, Shumway M, Zaborsky J, Feldblyum T, Subbu V, Spiro DJ, Sitz J, Koo H, Bolotov P et al. 2005. Large-scale sequencing of human influenza reveals the dynamic nature of viral genome evolution. Nature 437(7062): 1162–1166.

Hansen TH, Bouvier M. 2009. MHC class I antigen presentation: learning from viral evasion strategies. Nature reviews Immunology 9(7): 503–513.

Holland J, Spindler K, Horodyski F, Grabau E, Nichol S, VandePol S. 1982. Rapid evolution of RNA genomes. Science 215(4540): 1577–1585.

Holmes EC. 2011. What does virus evolution tell us about virus origins? Journal of virology 85(11): 5247. 5251.

Hu Y, Zandi R, Anavitarte A, Knobler CM, Gelbart WM. 2008. Packaging of a polymer by a viral capsid: the interplay between polymer length and capsid size. Biophysical journal 94(4): 1428–1436.

Iyer LM, Balaji S, Koonin EV, Aravind L. 2006. Evolutionary genomics of nucleo-cytoplasmic large DNA viruses. Virus research 117(1): 156–184.

Kay MA, Glorioso JC, Naldini L. 2001. Viral vectors for gene therapy: the art of turning infectious agents into vehicles of therapeutics. Nature medicine 7(1): 33–40.

Keese PK, Gibbs A. 1992. Origins of genes: “big bang”. or continuous creation? Proceedings of the National Academy of Sciences of the United States of America 89(20): 9489–9493.

Koonin EV. 2003. Comparative genomics, minimal gene-sets and the last universal common ancestor. Nature reviews Microbiology 1(2): 127–136.

Krakauer DC. 2000. Stability and evolution of overlapping genes. Evolution; international journal of organic evolution 54(3): 731–739.

Krakauer DC, Plotkin JB. 2002. Redundancy, antiredundancy, and the robustness of genomes. Proceedings of the National Academy of Sciences of the United States of America 99(3): 1405. 1409.

Lidmar J, Mirny L, Nelson DR. 2003. Virus shapes and buckling transitions in spherical shells. Physical review E, Statistical, nonlinear, and soft matter physics 68(5 Pt 1): 051910.

Liu H, Wu X, Newman M, Shaw GM, Hahn BH, Kappes JC. 1995. The Vif protein of human and simian immunodeficiency viruses is packaged into virions and associates with viral core structures. Journal of virology 69(12): 7630–7638.

Lynch M. 2010. Evolution of the mutation rate. Trends in genetics : TIG 26(8): 345–352.

Masson P, Hulo C, De Castro E, Bitter H, Gruenbaum L, Essioux L, Bougueleret L, Xenarios I, Le Mercier P. 2013. ViralZone: recent updates to the virus knowledge resource. Nucleic acids research 41(Database issue): D579–583.

Mizokami M, Orito E, Ohba K, Ikeo K, Lau JY, Gojobori T. 1997. Constrained evolution with respect to gene overlap of hepatitis B virus. Journal of molecular evolution 44 Suppl 1: S83–90.

Moreira D, Lopez-Garcia P. 2009. Ten reasons to exclude viruses from the tree of life. Nature reviews Microbiology 7(4): 306–311.

Moreno E, Ojosnegros S, Garcia-Arriaza J, Escarmis C, Domingo E, Perales C. 2014. Exploration of sequence space as the basis of viral RNA genome segmentation. Proceedings of the National Academy of Sciences of the United States of America 111(18): 6678–6683.

Noack J, Bernasconi R, Molinari M. 2014. How viruses hijack the ERAD tuning machinery. Journal of virology 88(18): 10272–10275.

Novella IS, Presloid JB, Taylor RT. 2014. RNA replication errors and the evolution of virus pathogenicity and virulence. Current opinion in virology 9: 143–147.

Pavesi A, De Iaco B, Granero MI, Porati A. 1997. On the informational content of overlapping genes in prokaryotic and eukaryotic viruses. Journal of molecular evolution 44(6): 625–631.

Pavesi A, Magiorkinis G, Karlin DG. 2013. Viral proteins originated de novo by overprinting can be identified by codon usage: application to the “gene nursery”. tof Deltaretroviruses. PLoS computational biology 9(8): e1003162.

Petrov AS, Harvey SC. 2008. Packaging double-helical DNA into viral capsids: structures, forces, and energetics. Biophysical journal 95(2): 497–502.

Ploegh HL. 1998. Viral strategies of immune evasion. Science 280(5361): 248–253.

Pruitt KD, Tatusova T, Brown GR, Maglott DR. 2012. NCBI Reference Sequences (RefSeq): current status, new features and genome annotation policy. Nucleic acids research 40(Database issue): D130– 135.

Rancurel C, Khosravi M, Dunker AK, Romero PR, Karlin D. 2009. Overlapping genes produce proteins with unusual sequence properties and offer insight into de novo protein creation. Journal of virology 83(20): 10719–10736.

Rappoport N, Linial M. 2012. Viral proteins acquired from a host converge to simplified domain architectures. PLoS computational biology 8(2): e1002364.

Rogozin IB, Spiridonov AN, Sorokin AV, Wolf YI, Jordan IK, Tatusov RL, Koonin EV. 2002. Purifying and directional selection in overlapping prokaryotic genes. Trends in genetics : TIG 18(5): 228–232.

Sabath N, Wagner A, Karlin D. 2012. Evolution of viral proteins originated de novo by overprinting. Molecular biology and evolution 29(12): 3767–3780.

Schneemann A, Schneider PA, Lamb RA, Lipkin WI. 1995. The remarkable coding strategy of borna disease virus: a new member of the nonsegmented negative strand RNA viruses. Virology 210(1): 1–8.

Shepherd CM, Borelli IA, Lander G, Natarajan P, Siddavanahalli V, Bajaj C, Johnson JE, Brooks CL, 3rd, Reddy VS. 2006. VIPERdb: a relational database for structural virology. Nucleic acids research 34(Database issue): D386–389.

Smith GJ, Vijaykrishna D, Bahl J, Lycett SJ, Worobey M, Pybus OG, Ma SK, Cheung CL, Raghwani J, Bhatt S et al. 2009. Origins and evolutionary genomics of the 2009 swine-origin H1N1 influenza A epidemic.Nature 459(7250): 1122–1125.

Snijder J, Uetrecht C, Rose RJ, Sanchez-Eugenia R, Marti GA, Agirre J, Guerin DM, Wuite GJ, Heck AJ, Roos WH. 2013. Probing the biophysical interplay between a viral genome and its capsid. Nature chemistry 5(6): 502–509.

UniProt C. 2014. Activities at the Universal Protein Resource (UniProt). Nucleic acids research 42(Database issue): D191–198.

Van Etten JL. 2003. Unusual life style of giant chlorella viruses. Annual review of genetics 37: 153–195.

Veeramachaneni V, Makalowski W, Galdzicki M, Sood R, Makalowska I. 2004. Mammalian overlappingenes: the comparative perspective. Genome research 14(2): 280–286.

Vignuzzi M, Stone JK, Arnold JJ, Cameron CE, Andino R. 2006. Quasispecies diversity determines pathogenesis through cooperative interactions in a viral population.Nature 439(7074): 344–348.

Worobey M, Holmes EC. 1999. Evolutionary aspects of recombination in RNA viruses. The Journal of general virology 80 (Pt 10): 2535–2543.

Wright JF. 2014. AAV empty capsids: for better or for worse? Molecular therapy : the journal of the American Society of Gene Therapy 22(1): 1–2.

Zandi R, van der Schoot P, Reguera D, Kegel W, Reiss H. 2006. Classical nucleation theory of virus capsids. Biophysical journal 90(6): 1939–1948.

